# Comparative transcriptomic insights into the evolutionary origin of the tetrapod double cone

**DOI:** 10.1101/2024.11.04.621990

**Authors:** Dario Tommasini, Takeshi Yoshimatsu, Tom Baden, Karthik Shekhar

## Abstract

The tetrapod double cone is a pair of tightly associated cones called the “principal” and the “accessory” member. It is found in amphibians, reptiles, and birds, as well as monotreme and marsupial mammals but is absent in fish and eutherian mammals. To explore the potential evolutionary origins of the double cone, we analyzed single-cell and -nucleus transcriptomic atlases of photoreceptors from six vertebrate species: zebrafish, chicken, lizard, opossum, ground squirrel, and human. Computational analyses separated the principal and accessory members in chicken and lizard, identifying molecular signatures distinguishing either member from single cones and rods in the same species. Comparative transcriptomic analyses suggest that both the principal and accessory originated from ancestral red cones. Furthermore, the gene expression variation among cone subtypes mirrors their spectral order (red *→* green *→* blue *→* UV), suggesting a constraint in their order of emergence during evolution. Finally, we find that rods are equally dissimilar to all cone types, suggesting that they emerged before the spectral diversification of cones.

## 1 Results

Eight types of photoreceptors have been identified across the retinas of extant vertebrates: rods, four types of single cones (red, green, blue, UV), “blue rods”, and the two members of the double cones (DCs) ^1^. Rods and single cones are ubiquitous across vertebrates, and therefore were likely present in the retina of the common vertebrate ancestor *∽* 500 million years ago (MYA) ^2–4^. In contrast, DCs and blue rods likely emerged later (*∽*390 MYA), around the beginning of vertebrate life on land ^1^. The DC is so called because it is composed of two tightly associated photoreceptors, the principal (DC-P) and the accessory (DC-A) ^5,6^. DCs can be quite numerous, comprising up to 40% of the cones in some avian retinas ^7^. Like ancestral red cones, DCs express the long-wavelength sensitive (LWS) opsin and are, therefore, expected to be broadly tuned for long wavelengths ^8,9^. However, their larger size and absence in the fovea of some raptorial birds suggests they may support fast achromatic processing ^10,11^. Additionally, DCs have been proposed to mediate light-dependent magnetoreception and/or polarized light sensing because they express the magnetically sensitive protein cryptochrome 4^12^ and form highly ordered arrays ^13,14^. However, to our knowledge, outside of salamanders ^8,9^, DCs have never been directly recorded from in an intact retina.

The presence of DCs in birds, reptiles, amphibians, monotremes, and marsupials, but not in fish and other mammals suggests that they arose in the common ancestor of tetrapods and were later lost in eutherian mammals ^1,4^. While their evolutionary origin is debated, the phylogeny of DCs (**Figure 1A**) suggests they may have evolved from ancestral vertebrate single cones. One hypothesis – based on morphological similarity to pairs of red and green cones in fish retinas ^15^ – is that DC-P evolved from the ancestral red cone and DC-A evolved from the ancestral green cone ^1^. Alternatively, because both DC-P and DC-A express the LWS opsin, it has been suggested that both DC members may have evolved from the ancestral red cone ^8,10^. However, neither of these hypotheses has been tested rigorously.

**Figure 1.**
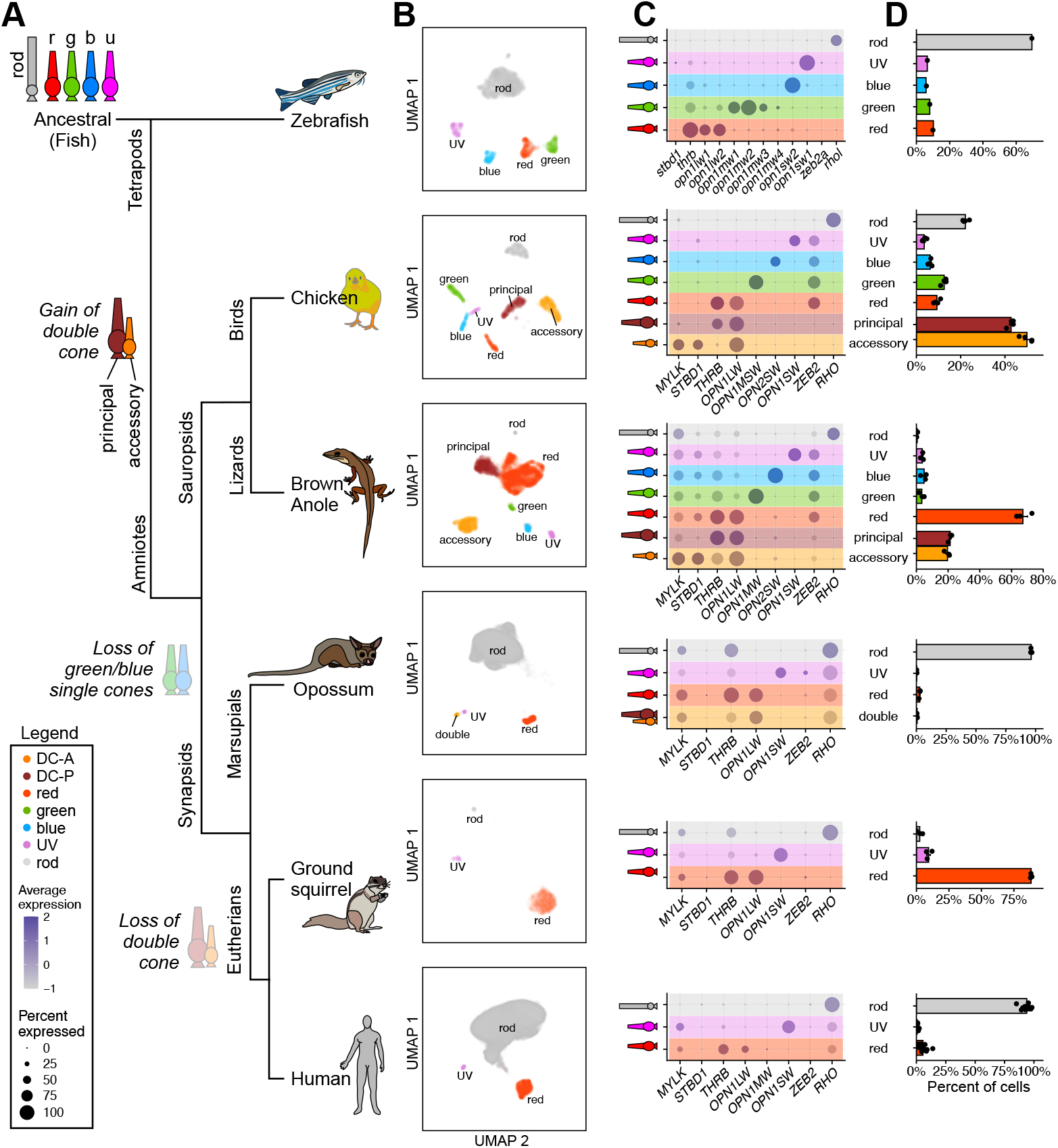
Single-cell atlases of photoreceptors across six vertebrate species. A) Phylogenetic tree showing the putative evolutionary history of photoreceptor types in the six studied vertebrates. B) 2D Uniform Manifold Approximation (UMAP) ^21^ embeddings of single-cell atlases. Each subpanel (row) represents data from a different species. Colors correspond to photoreceptor types as in panel A. C) Dot plots showing expression of marker genes corresponding to panel B. Color shows average expression scaled for each gene across photoreceptor types. Circle size corresponds to the percentage of cells in the cluster that express that gene. See the legend on bottom left. D) Relative proportion of photoreceptor types. Points correspond to biological replicates. The original chicken atlas had a third cluster composed of DC-P and DC-A doublets, which is not shown here (see **Figure S1**).

### 1.1 Comparative analysis of photoreceptor atlases

We addressed this question through a comparative transcriptomics approach, hypothesizing that similarity in gene expression might reveal the evolutionary relationships between vertebrate ciliary photoreceptor types. To this end, we analyzed published single-cell (sc) and single-nucleus (sn) RNA-sequencing (RNA-seq) atlases from six vertebrate species: zebrafish (*D. rerio*) ^16^, chicken (*G. gallus*) ^17^, brown anole lizard (*A. sagrei*) ^18^, opossum (*M. domestica*) ^18^, thirteen-lined ground squirrel (*I. tridecemlineatus*) ^18^, and human (*H. sapiens*) ^19^ (**Figure 1**). By applying dimensionality reduction and clustering to each species atlas, we identified transcriptomic clusters corresponding to rods, single cones, and putative double cones (**Figure 1B, Table S1**). As expected, zebrafish, chicken, and lizard contained rods and the full complement of single cones (red, green, blue, UV), while opossum, squirrel, and human contained rods and only two types of single cones (red and UV). Note that we annotated mammalian cones based on their ancestry rather than spectral sensitivity (see **Methods**). For instance, although humans possess both a red and a green cone, these are both derived from the ancestral red cone, resulting in nearly identical transcriptomic profiles ^20^. While the two types of cones have been separated using supervised approaches elsewhere ^20^, we choose to retain them as a single cluster in this study. Furthermore, the human blue cone is derived from the ancestral UV cone and expresses a blue-shifted UV opsin ^4^. Similarly, squirrel green cones express a green-shifted red opsin, and are thought to have derived from red cones ^4^.

### 1.2 Molecular identification of double cones

Beyond rods and single cones, we found two additional clusters in chicken and lizard representing putative double cone members (**Figure 1B**). These two clusters expressed *OPN1LW*, the gene encoding the LWS opsin, but unlike the single cones in chicken and lizard, they did not express *ZEB2* (**Figure 1C**). In addition, these clusters were present in approximately a 1:1 stoichiometric ratio in chicken and lizard (**Figure 1D**). One of these clusters was enriched for the red cone marker *THRB* ^22,23^, while the other selectively expressed *STBD1*. We hypothesize that the *OPN1LW*^+^*THRB*^+^*STBD1*^*−*^ cluster represents DC-P and the *OPN1LW*^+^*THRB*^*−*^*STBD1*^+^ cluster represents DC-A (see **Methods**), and refer to these as DC-P and DC-A respectively hereafter. We note that, unlike chicken and lizard, fish do not contain additional clusters corresponding to DCs (also, see **Discussion**).

We performed fluorescence *in situ* hybridization chain reaction (HCR) ^24^ for *OPN1LW* and *STBD1* in anole lizard retinal sections, which confirmed that the relative abundance of photoreceptors with these expression patterns are consistent with the sc/snRNA-seq data (**Figure S2A-D**). In opossum, we identified a cluster separate from the red cone cluster that was *OPN1LW*^+^*ZEB2*^*−*^ and expressed low levels of *THRB*, suggesting that these cells may represent DCs, but there were not enough cells to resolve DC-P and DC-A separately (**Figure 1B**). As expected, zebrafish and the two eutherian mammals (squirrel and human) each contained a single *OPN1LW+* cluster corresponding to red single cones. Taken together, these results are consistent with the notion that double cones evolved with or after the first emergence of vertebrate life on land and were later lost in eutherian mammals ^1^.

The relative abundances of photoreceptor types within each species were consistent across biological replicates and were comparable to previous reports ^7,25–27^ (**Figures 1D, S2E**,**F**). Double cones are the most prevalent photoreceptor type in chicken (*∽*45%) and the second-most prevalent type in lizard (*∽*20%), but were quite rare in opossums (*<*1%). Rod frequencies exhibit the highest variation, being *>*90% in the nocturnal opossum and *<*1% (but not absent) in the diurnal lizard.

### 1.3 Correspondence between ancestral photoreceptors

Before investigating the evolutionary origin of double cones, we first asked whether we could recover the long-suspected evolutionary relationships of the ancestral photoreceptors, i.e. single cones and rods. We began with an integration approach that relies on 1:1 orthologous genes ^18,28^, but this method failed to fully separate the different cone types (**Figure 2A**). Therefore, we applied an alternative approach, SAMap, which incorporates complex gene homology relationships and iterative refinement ^29^. SAMap recovered the expected photoreceptor homologies among zebrafish, chicken, and lizard photoreceptors, each of which have the full complement of single cones and rods. The mammalian photoreceptor types correctly co-clustered with their non-mammalian counterparts (**Figure 2C,D**). The conserved transcriptional signatures for the five ancestral photoreceptors included their respective opsins -*RHO* in rods, *OPN1LW* in red cones, *OPN1MW* in green cones, *OPN2SW* in blue cones, and *OPN1SW* in UV cones -as well as known photoreceptor genes such as *GNGT1, PDE6B, PDE6H*, and *THRB* ^30,31^, demonstrating SAMap’s ability to correctly identify functional orthologs (**Figure 2D, S3**). Additionally, SAMap also identified several paralogous relationships among genes that may represent duplication events – as is the case for several genes in zebrafish (**Figure S3**), which has an additional whole genome duplication relative to other vertebrates ^32^.

**Figure 2.**
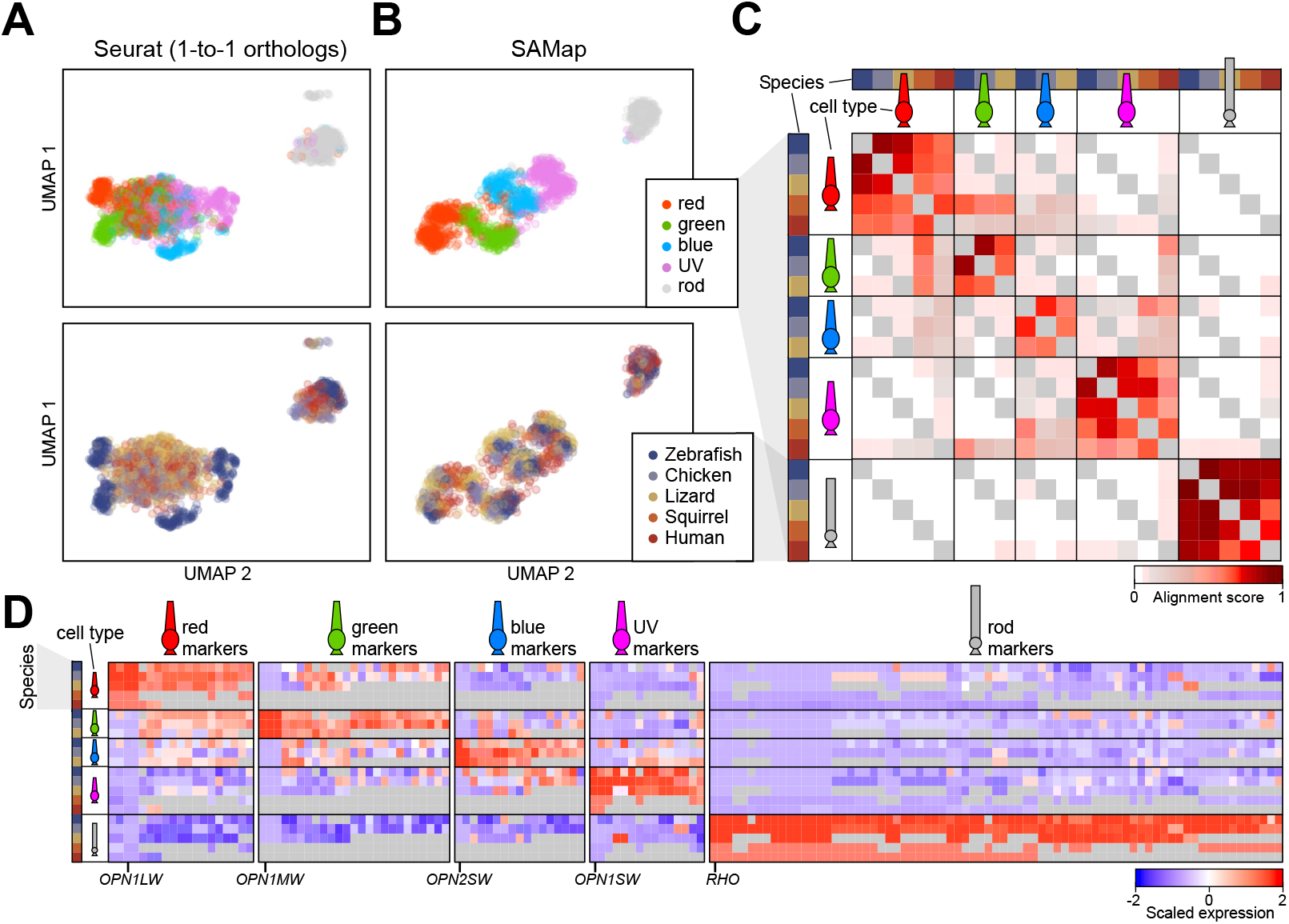
Cross-species integration of ancestral photoreceptors. A) 2D visualization of single cone types and rods from six species integrated using Seurat v4^28^, resulting in the intermixing of cone types. B) Same as panel A, but with integration performed using SAMap ^29^, which preserves cone type identity. C) Heatmap of SAMap alignment scores for photoreceptor types. Alignment scores represent averaged similarity in the k-nearest neighbor graph between two clusters (see **Methods**). Evolutionarily related photoreceptors have high alignment scores. D) Heatmap of scaled gene expression showing the top conserved transcriptional signatures of ancestral photoreceptor types. Since genes can have multiple homologs, each column represents a unique combination of homologous genes across species. Gray values indicate that no gene homolog was found for that species. Only key genes are highlighted; for all gene names, see **Figure S3**.

### 1.4 Molecular relationship of double cone members to single cones

Having confirmed the validity of the cross-species mapping approach for single cones and rods, we next analyzed the transcriptional relationships between the DCs and the single cones. We hypothesized that each member would exhibit the highest transcriptional similarity to the single cone with whom it shares a common ancestor. We first used SAMap to compare the chicken and lizard atlases. All photoreceptor types, including DC-P and DC-A, aligned in a 1-to-1 fashion (**Figure 3A**), indicating that the double cone members of chicken and lizard are transcriptionally homologous. Interestingly, DC-P exhibited a weak affinity to the red cone, foreshadowing the results presented below.

**Figure 3.**
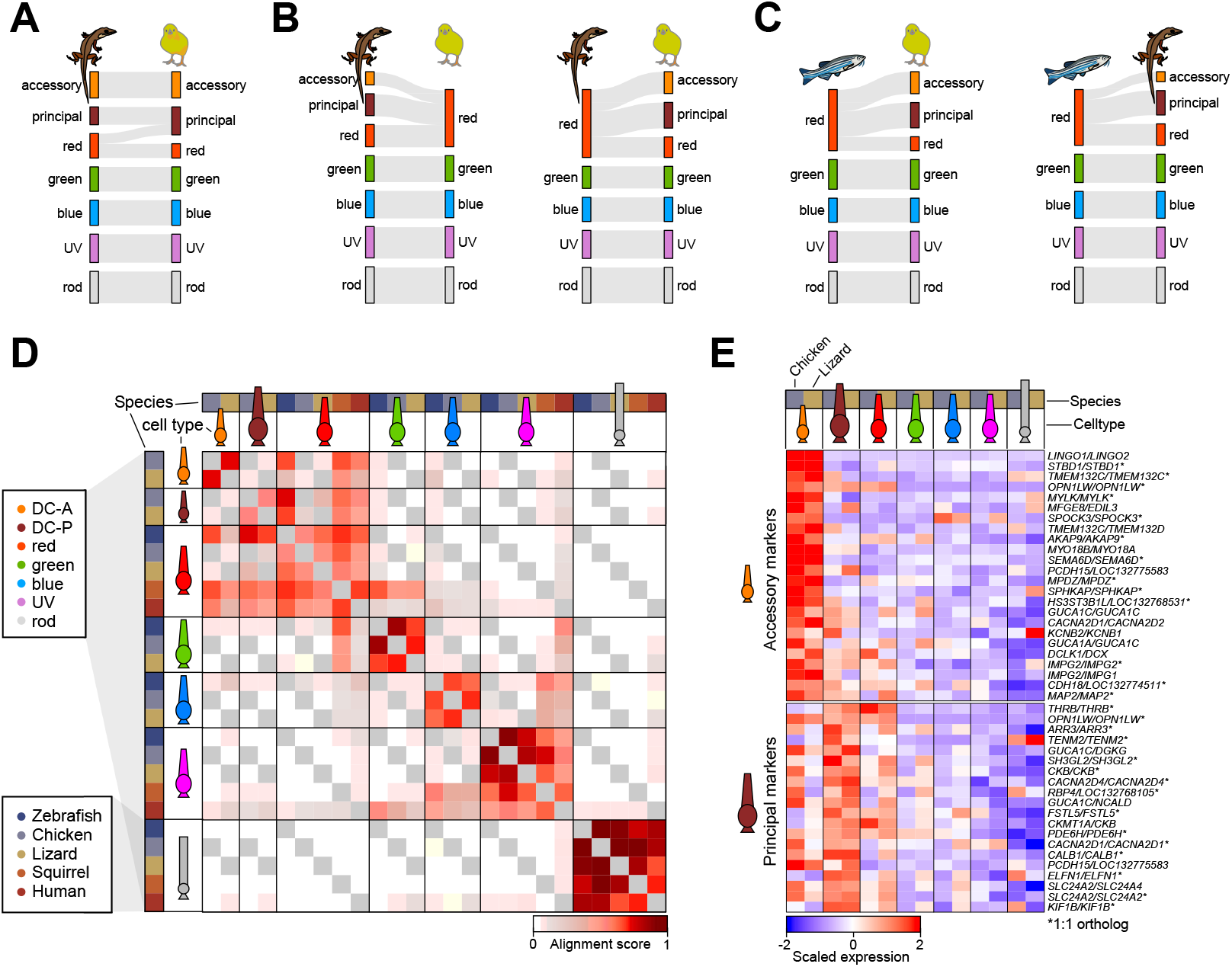
Transcriptomic similarity of DC-P and DC-A with the red cone suggests a shared evolutionary history. A) Sankey diagram showing SAMap alignment scores between chicken and lizard when aligning all photoreceptors. B) Like A, but with the removal of chicken double cones (*left*) or lizard double cones (*right*). C) Alignment of chicken with zebrafish (*left*) and lizard with zebrafish (*right*). D) Heatmap of SAMap alignment scores when aligning photoreceptors across zebrafish, chicken, lizard, squirrel, and human. E) Top conserved genes of DC-A and DC-P, shown as a heatmap of scaled gene expression.

Next, we conducted a series of computational experiments to determine the closest ancestral photoreceptor counterpart(s) of the DC members. To begin, we repeated the chicken/lizard mapping, but this time, we removed the DCs from one species – either chicken or lizard. Although our method allows multi-mapping between clusters, we found that in either case DC-P and DC-A each mapped specifically to the red single cone in the other species (**Figure 3B**). Next, we aligned chicken and lizard atlases to the zebrafish atlas, which lacks DCs (**Figure 3C**); again, both DC members mapped to the zebrafish red cone. Finally, we aligned all five species, including squirrel and human (**Figure 3D**). All the photoreceptor types aligned nearly exclusively to each other except red cones, DC-P and DC-A, which formed a block of high alignment scores with each other. Sporadic instances of cross-mappings were observed between red cones and green cones and between blue cones and UV cones, although these were consistently weaker than the DC-P *→* red and DC-A *→* red mappings. Notably, rods exhibited no cross-mappings with cone types and were the most distinct photoreceptor type.

We also analyzed within-species cell type similarity by calculating the expression correlation among all photoreceptor types (**Figure S4A**). In both chicken and lizard, DC-P was transcriptionally most similar to the red cone. On the other hand, DC-A was most similar to DC-P, followed by the red cone. Together, these findings indicate that both DC members are most transcriptionally similar to the red cone out of all the ancestral photoreceptors. Correlations with the green cone had consistently lower values, followed by blue and UV cones, and then rods. The expression correlation values were also inversely proportional to the number of differentially expressed genes (DEGs) among types (**Figure S4B**).

As shown in **Figure 3E**, DC-P is characterized by high expression of *THRB, FSTL5, ARR3, ELFN1*, and *PDE6H*, a signature that is highly similar to that of red cones (**Figure 3E**). DC-A has a more distinct signature (**Figure 3A**) and is marked by the conserved expression of *STBD1, MYLK*, and *TMEM132C* (**Figure 3E**). To further investigate the molecular characteristics of DCs, we conducted differential expression tests to identify markers that distinguish the two DC members from each other and the DC members from red cones within each species (**Figure S5**). We found several conserved signatures, reported in **Table S2**. For instance, both chicken and lizard DC-P are *SOX5+/SPOCK3-* while both DC-A are *SOX5-/SPOCK3+*. However, only one gene, *ZEB2*, distinguishes DCs from red cones in both species (**Figure S5** and **Table S2**).

### 1.5 Cone gene expression mirrors their spectral order

In zebrafish, chicken, and lizard, we observed that the transcriptional relationships between photoreceptor types mirror their spectral arrangement: blue cones are most similar to UV cones and green cones, while green cones are most similar to blue and red cones (**Figure S4**). Using principal component analysis (PCA), we confirmed that the second principal component (PC2) reliably captures the spectral order of the various cone types (**Figure 4A**). The genes driving this principal component in zebrafish were reproducible in two separate datasets ^16,31^ (**Figure S6**). Furthermore, the PCA revealed that DC-P and DC-A are most proximal to the red cone and lie in the “infrared” region of PC2 (**Figure 4A**).

**Figure 4.**
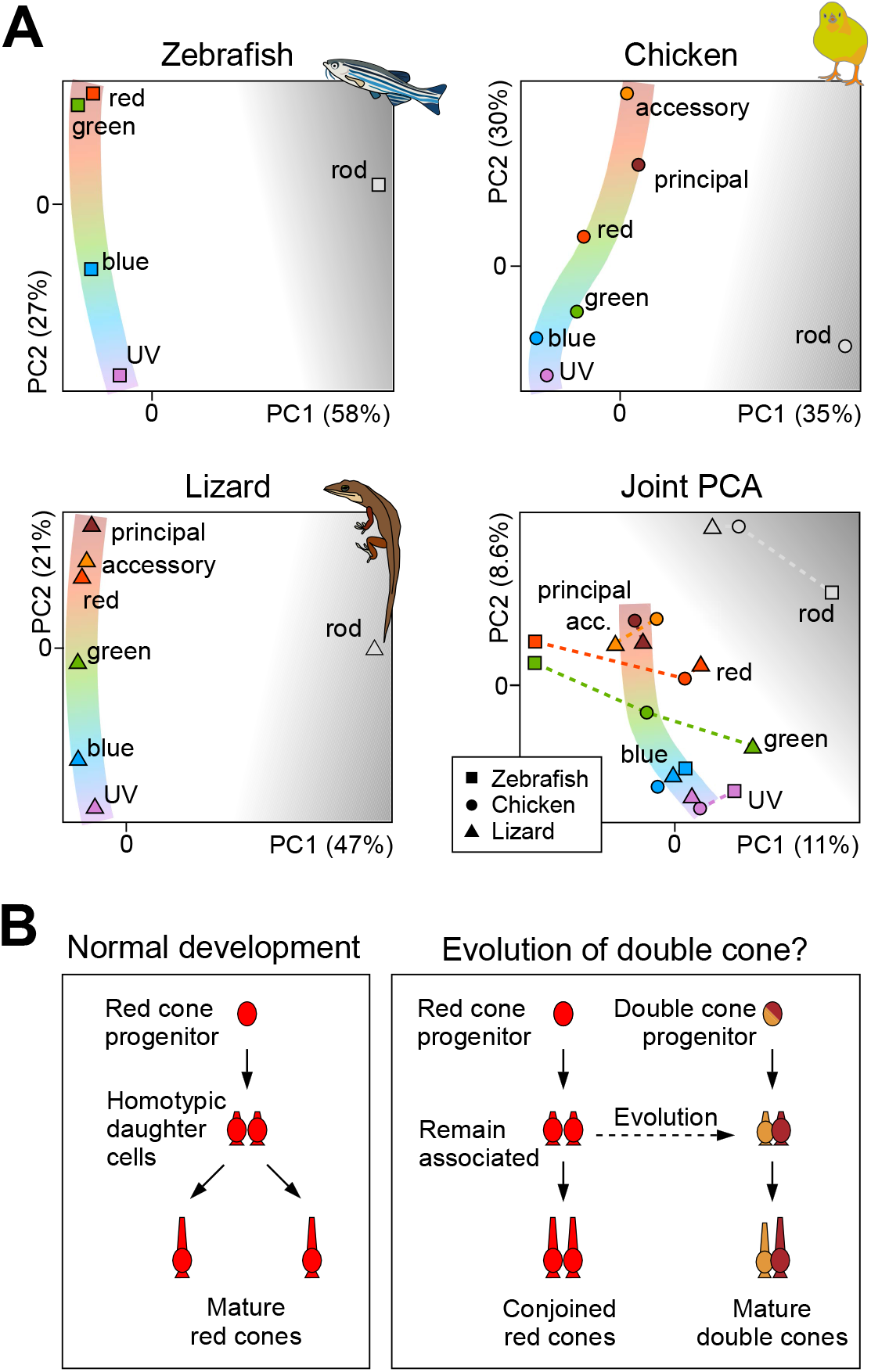
Transcriptomic variation among photoreceptors mirrors their spectral relationships. A) PCA embedding of photoreceptors in zebrafish, chicken, and lizard separately and in the joint expression space derived from SAMap. To reduce species-specific noise, we binarized the scaled expression values as 0 (not expressed) or 1 (expressed) based on a set threshold (see **Methods**). Photoreceptors lie on a gene expression manifold that mirrors the color spectrum. B) Proposed model of how double cones may have arisen in from ancestral red cones. *Left panel:* Experiments in zebrafish ^33^ indicate that single cones arise during development via symmetric terminal divisions of committed precursors ^34^ *Right panel:* In an ancestral tetrapod, a mutated red-cone precursor gave rise to daughter cells expressing the LWS opsin that remained associated as they mature. Over evolutionary timescales these associated daughter cells acquired distinct properties, becoming the DC-P and the DC-A.

### 1.6 Rods are equally dissimilar to all cones

Rods have long been speculated to have evolved from cones ^2,35^. The protein sequence similarity between rhodopsin and the green-sensitive opsin has been used as evidence to suggest that rods evolved from green cones ^2,36,37^, whereas molecular signatures suggest a UV cone origin ^38,39^. As an attempt to distinguish between these scenarios, we examined the transcriptional similarity of rods and different cone types. We found that rods are highly dissimilar to all cone types and have no consistent affinity towards any cone type across zebrafish, chicken, and lizard (**Figure S4**). In all PCAs, the first PC, which captures the highest variance, always separated rods and cones (**Figure 4A**). Moreover, SAMap alignment scores between rods of different species were high (*>*0.6), but alignment scores between rods and cone types were negligible (*<*0.05) (**Figures 2C, 3D**). These results suggest that rods may not be evolutionarily derived from any specific cone type. Instead, it suggests that ancestral rods diverged from ancestral cones prior to the spectral diversification of cones.

## 2 Discussion

In this study, we integrated single-cell atlases across vertebrate species that span the emergence and loss of DCs in the tetrapod lineage. We provide molecular markers for the principal and accessory members of the tetrapod double cone and, through comparative analyses, suggest they evolved from the red cone. Our main contributions are as follows:

### Transcriptomic atlases of photoreceptors

Through clustering analyses, we identified rods and single cones in each species and pinpointed clusters corresponding to DCs in opossum, lizard, and chicken. We isolated the principal and accessory member of the DC in chicken and lizard, but we were unable to do so in the opossum because of their low abundance. Cross-species orthology among photoreceptor types has been long-posited based on opsin expression alone (**Figure 2C**). Our whole-transcriptome comparisons corroborate these orthologies, and, moreover, furnish additional markers reported that may help verify the molecular identity of photoreceptors in other species (**Figure 2D, S3, 3E**).

Comparisons of cell type composition across species show that while DCs are highly abundant in birds, they are less frequent in reptiles, and quite rare in marsupials (**Figure 1D**). In all six species, the relative abundance of red and green cones is higher than the blue and UV cones. Furthermore, while the classical literature has largely regarded diurnal reptiles as rodless ^40,41^, we find that anole lizards have a very small population of rods (**Figure 1B-D**). The wide variations in photoreceptor composition likely reflect the differences in the diverse ecological habitats of these species and their distinct evolutionary histories ^42^.

### Evolutionary origin and molecular identity of the DC

Our integrative analyses suggest that both DC-P and DC-A arose from a duplication of ancestral red cones. This conclusion is based on the higher degree of transcriptomic similarity between DC-P and DC-A with the extant red cone compared to the other single cone types. Furthermore, the experimental observation that zebrafish cones arise from symmetric terminal divisions of dedicated precursors ^33,34,43^ suggests the following scenario for the genesis of DCs: during the beginning of vertebrate life on land, a mutated version of the red cone precursor arose, whose daughter cells did not separate at birth. Over evolutionary time scales, these two daughter cells developing distinct characteristics and became the modern-day principal and accessory members of the double cone (**Figure 4B**). Developmental scenarios that produce a conjoined red/green cone seem unlikely, as asymmetric terminal divisions have, to our knowledge, not been observed in fish. We provide genes that distinguish the DC members from each other and from cones in lizard and chicken (**Figures 3E, S5** and **Table S2**). In addition to enabling DCs to be labeled and manipulated experimentally, these genes may be useful in understanding their function (**Figures 3E, S5**). Compared to DC-P, DC-A is more distinct from the red cone (**Figure S4**). Genes enriched in DC-A include those involved in calcium-dependent cell-cell adhesion (*PCDH15, CDH18*), protein kinase A activity (*SPHKAP, AKAP9*), and myosin-related proteins that are highly expressed in muscle cells (*MYLK, MYO18A/B, MAP2, STBD1*). The expression of muscle-related genes may reflect high metabolic demands and/or the need to traffic molecules (e.g. retinal) intracellularly at high rates. A candidate of particular interest is *STBD1*, which co-localizes with glycogen stores and is thought to bind glycogen and anchor it to membranes ^44^. Given that *STBD1* is expressed specifically in DC-A and not in other photoreceptors (**Figure 1C**), it may be associated with the enlarged paraboloid (glycogen-containing organelle) that is unique to DC-A ^13,41,45^. This may point to the DC-A serving a metabolic function in addition to its visual role. We note that our annotation of the DC clusters as DC-P and DC-A is tentative and remains to be fully validated.

### Fish “double cones”

One long-standing confusion concerns the putative presence of “double cones” in the retinas of teleost fish, such as zebrafish. Like many surface-dwelling and diurnal fish, adult zebrafish have a crystalline photoreceptor mosaic with fixed 2:2:1:1 (R:G:B:U) cone stoichiometry. These are arranged into alternating rows of R/G and B/U cones, with R/G cones arranged into intimate pairs, often with gap junctions in between them ^15^. This anatomical arrangement of the R/G pairs has long been taken as a signature of double cones, incorrectly linking them to the “true” double cones of tetrapods. However, teleost R/G pairs and tetrapod double cones are distinct sets of photoreceptors ^46^.

Perhaps the simplest way to ascertain this may be during early development: fish retinas grow throughout life, and in their early larval stages, the crystalline mosaic is not yet formed, despite the spatially independent presence of all four single cone types in the eyes of many larval fish ^15^. Over developmental time, this “larval patch” gradually transitions towards the adult mosaic, such that the transition from unpaired to paired R/G single cones becomes “inscribed” into the retinal anatomy of adult fish (i.e. with the larval patch being located near the optic disc). For further discussion, see ^46^. The independent transcriptomic clusters formed by tetrapod single and double cones found in all non-eutherian tetrapods tested (e.g. **Figure 1**) further cement this notion.

### Molecular relationships among photoreceptor types

Molecular similarities across the four ancestral single cones add further credence to the prevailing notion that all cones share a common origin. Within this group, we find that molecular similarities of cones across species consistently follow their spectral order: red *→* green *→* blue *→* UV. This relationship suggests that spectrally neighboring cones are evolutionarily related, which may provide a potential explanation for “spectral block wiring” – the finding that spectrally neighboring cones have an above-chance tendency to co-wire into postsynaptic circuits ^4,45,47–49^. Alternatively, though arguably less plausibly, cones with shared spectral sensitivity may have acquired similar gene expression patterns through convergent evolution. Finally, it remains unclear which cone type came first. Their spectral (and molecular) order hints that it could either be the red or UV cone. The earliest photoreceptors, long predating eyes, were likely UV-sensitive since retinal by itself (with no opsin) is UV-sensitive ^50^. On the other hand, the red cone is considered to be the most functionally critical photoreceptor ^4,51^, and the pineal organ also has red-cone-like receptors ^52^. Thus, arguments can be made for both the red and the UV cones emerging first.

Lastly, we find no molecular evidence that rods are related to green cones (**Figure 3D, 4A, S4**), as the protein sequence similarity of opsins might suggest ^2,36,37^. It is possible that the rod simply diverged beyond molecular recognition from its hypothetical cone predecessor. However, the fact that rods are equally dissimilar to all cone types suggests that rods diverged prior to the spectral diversification of cones.

## 3 Methods

### 3.1 Choice of species

To investigate the evolution of DCs, we needed to sample 1) species that diverged before the emergence of DCs, 2) species with DCs, and 3) species that have since lost DCs:

For 1), we used zebrafish, which possesses the ancestral photoreceptor complement (rods plus red, green, blue, and UV cones). We attempted to include photoreceptors from goldfish (another teleost fish) ^53^, but we were unable to identify blue and UV cones, possibly due to low sampling and/or poor opsin annotation.

For 2), we used chicken, lizard, and opossum, which contain DCs ^4^. We attempted to include the amphibian *Xenopus* ^54^, which also has DCs. Although we were able to identify several red cone clusters in *Xenopus* – indicating the likely presence of red cones and DCs – the overall cell count was too low for a comprehensive annotation.

For 3), we used two eutherian mammals: human ^19^ and squirrel ^18^. We had first attempted to use mouse. However, some mouse cones are known to co-express both *OPN1MW* and *OPN1SW* along a dorsoventral gradient with *OPN1SW* enriched in the ventral retina ^42,55^. Due to this, we were unable to identify discrete populations of single-opsin cones (i.e. *OPN1MW+* or *OPN1SW+*) across multiple mouse atlases ^56,57^, instead observing a single cluster with graded opsin expression. Thus, we decided to use squirrel, in which we could easily distinguish the two cone clusters (**Figure 1B**).

### 3.2 Alignment of sc/snRNA-seq data

We retrieved pre-processed count matrices from published studies for zebrafish ^16,31^, chicken ^17^, brown anole lizard ^18^, opossum ^18^, squirrel ^18^, and human ^19^. However, we noticed that the lizard transcriptome did not contain annotations corresponding to the green and the blue opsin. Therefore, we re-aligned the raw sequencing data from lizard ^18^ to a newer transcriptome assembly, *Anolis sagrei v2*.*1*, available on NCBI (https://www.ncbi.nlm.nih.gov/datasets/-genome/GCF_025583915.1/). **Table S1** lists the gene names corresponding to opsins in each species. As we could not identify DC components in opossum, we also attempted to re-align the opossum raw data to a newer NCBI assembly (mMonDom1.pri 2023) but this yielded similar results to the original ENSEMBL-aligned data ^18^. This suggests that more opossum cells be needed to resolve DC-P and DC-A.

### 3.3 Annotation of cell atlases

To ensure high-quality annotations, we applied a consistent pipeline to assemble the photoreceptor atlas for each species. This involved reclustering the original dataset, filtering doublets, and annotating clusters based on the expression of opsin genes (**Table S1**). Clusters were annotated as rods, single cones (red, green, blue, UV) and putative double cone principal and accessory members. As noted in the main text, mammalian cone types were annotated based on their ancestry. Thus, the human red and green cones, which are both derived from ancestral red cones, were annotated together as “red cones” ; human blue cones, derived from the ancestral UV cone, was annotated “UV cones”; and squirrel green cones, derived from ancestral red cones, were annotated as “red cones” (**Figures 1B,C**).

For zebrafish, Ogawa and Corbo ^31^ described an additional cone cluster defined by the coexpression of *opn1mw4* and *opn1lw1* opsin genes (*opn1mw4/opn1lw1+* cones) in their scRNA-seq data. The authors hypothesized that this cluster might represent a unique subpopulation within the commonly observed R/G cone pairs in teleosts (see **Discussion**). Ogawa and Corbo also noted the graded expression of *opn1lw1/2* and *opn1mw1/2/3/4* in the red and green cones, consistent with the presence of region-specific subpopulations.

Encouragingly, we were able to reproduce Ogawa and Corbo’s observations in the adult zebrafish snRNA-seq dataset of Lyu et al. ^16^ (**Figures S7**). However, we could not detect *opn1mw4/opn1lw1+* cones in larval zebrafish at 4/5 days post fertilization, suggesting that *opn1mw4/ opn1lw1+* cones may arise at later developmental stages (**Figures S7A,B**). Finally, we note that the appearance of *opn1mw4/opn1lw1+* cones as a single cluster in both scRNA-seq and snRNA-seq datasets suggests that they represent a subpopulation of single cones rather than true double cones.

### 3.4 Identification of intact double cones

In their scRNA-seq atlas of the chicken retina, Yamagata et al. ^17^ identified three putative double cone clusters - DCa, DCb, and DCc. We hypothesized that one of these may represent intact DCs, with the other two being dissociated DC-P and DC-A cells. To test this, we used linear regression to fit the expression vector of each cluster based on the other two clusters (for e.g., *DCb ≈ αDCa* + *βDCc* + *γ*), where *DCi* is the gene expression vector (*i* = *a, b, c*), *α* and *β* are regression coefficients, and *γ* is the bias. We found that *DCb ≈* 0.5*DCa* + 0.56*DCc* (*p <* 10^*−*3^, **Figure S1C**), suggesting that DCb represents the intact double cone and DCa and DCb are its members. Consistent with this result, modeling either DCa or DCc as a linear combination of the other two clusters yielded subtractive combinations (**Figure S1C**), again consistent with DCb representing full double cones. Thus, we reannotated DCb as full intact DCs and omitted it from downstream analysis. Notably, clusters corresponding to intact DCs were found only in scRNA-seq data not snRNA-seq data. We used chicken as the basis for annotation of the less-studied anole lizard. We hypothesized that the DC cluster expressing the red cone marker *THRB* represents DC-P because of the morphological similarity between DC-P and red cones, leaving the *THRB*^*−*^*STBD1*^+^*MYLK*^+^ cluster as the putative DC-A.

### 3.5 Cross-species photoreceptor alignment using SAMap

SAMap analysis was run as follows. 1) An initial SAMap object was instantiated and the resulting BLAST homology graph was saved for later use, as this is the slowest step. 2) h5ad count files exported from Seurat were preprocessed using SAM ^58^ with 100 PCs, k=20 nearest neighbors, and 3000 genes. 3) SAMap was run pairwise for 3 iterations, with computing neighborhoods from keys set to true. 4) Alignment scores (average scores from the kNN graph) were extracted using the `get_mapping_scores` function in SAMap. We also used the function `GenePairFinder` with default parameters to identify conserved gene markers for photoreceptors in **Figure 2D** and **Figure 3E**.

Due to the large differences in photoreceptor type frequency across species, we downsampled each photoreceptor type within each species to 100 cells. This inherently introduces randomness in the mapping, so we repeated SAMap experiments 10 times with different random samples and took the median alignment scores across the 10 runs. This ensured that our results were robust and reproducible across different random samples of cells.

### 3.6 Differential gene expression analysis

For identifying differentially expressed genes (DEGs), we used the R package presto. For constructing the hierarchical clustering trees in **Figure S4B**, we used an average log2 fold change cutoff of 1 and adjusted p-value cutoff of 0.001. For exploration of DEGs distinguishing DCs from each other and from red cones (**Figure S5, Table S2**), we used more permissive cutoffs (average log2 fold change cutoff of 0.25 and adjusted p-value cutoff of 0.01).

### 3.7 Hierarchical clustering and principal component analysis (PCA)

Hierarchical clustering trees based on Pearson correlations (**Figure S4A**) were constructed using the pseudobulked expression of the top 2000 highly variable genes. Briefly, normalized counts were averaged in non-log space and then log-transformed using the log1p function in R. Correlations were computed across different single-cell clusters using the cor function in R.

For the PCAs within species (**Figure 4A**), we used the pseudobulked expression of the top 2000 highly variable genes to compute principal components (PCs). Normally, PCA is preceded by scaling across features. We observed that when computing PCs with centered and scaled expression data, the resulting PCs were dominated by contributions from noisy, lowly expressed genes. We found two strategies to avoid this noise. 1) We centered the data but did not scale prior to PCA. This effectively gives highly expressed genes more importance in the reduced dimension. 2) We require genes to be expressed above a certain threshold in one cluster (e.g. 5-15% of cells). These two methods led to very similar results, so we used centering without scaling since it did not require arbitrary hyperparameters, and was faster.

For the joint PCA in **Figure 4A**, we used the homology graph from SAMap ^29^ to transform the gene expression values of zebrafish and lizard into the chicken gene expression space. Briefly, the homology graph **A** is an *m*1 *× m*2 matrix, where *m*1 and *m*2 are the number of genes from species 1 and species 2 respectively, and whose entries denote the similarity between genes across species. These similarities are initialized from protein BLAST similarity, but then refined based on expression similarity ^29^. If 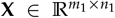 is the gene-by-type expression matrix for species 1 with *n*1 types, then 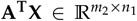 is the transformed gene-by-type matrix in the expression space of species 2. We scaled within each species to remove extensive batch effects across species, then subsetted to genes that are present across all three species (*∽*8000 genes), and horizontally concatenated the three gene-by-type matrices. To mitigate species-specific noise, we binarized these scaled expression values by setting genes whose expression was above an arbitrary threshold to 1 and the rest to 0. We used a threshold of -0.15, but other values around 0 gave similar results. We then ran PCA on the resulting binarized gene-by-type matrix.

### 3.8 In situ HCR and light microscopy

All procedures were performed in accordance with the Washington University in St. Louis, Institutional Animal Care and Use Committee (IACUC) guidelines. For *in situ* HCR, we used female adult green anoles purchased from Carolina Biological Supply, age unknown. Adult lizards were humanely euthanized with a high dose of anesthesia (Alfaxalone 60 mg/kg) subcutaneous injection. After decapitation, retinal tissues were dissected from the enucleated whole eyes by removing cornea, lens and epithelial layer in 1x PBS. The tissues were immediately fixed in 4% paraformaldehyde (Agar Scientific, AGR1026) in PBS for 20 min at room temperature, followed by three washes in PBS. The tissues were then sliced at 200 μm thickness using a tissue chopper. The standard *in situ* HCR was performed according to the manufacturer’s protocol using HCR Probe hybridization buffer, Probe Wash buffer, and Amplification buffer (Molecular Instruments). HCR probe sets and Amplifiers were custom-designed (**Table S3**). Hoechst 33342 was added to visualize nuclei during the wash step after the amplification step.

Confocal image stacks were taken immediately after the *in situ* HCR on a FV1000 microscope (Olympus) with a 40x oil immersion objective (HC PL APO CS2, Leica). Typical voxel size was 0.62 μm and 0.5 μm in the *x*-*y* and *z*, respectively. Contrast, brightness and pseudo-colour were adjusted for display in Fiji ^59^. Puncta were detected by thresholding the image stacks followed by 3D Object Counter in Fiji. For *OPN1LW*, nuclei with puncta signal were counted as positive. Briefly, we used Cellpose ^60^ to segment the nuclei, and then used a script from the 10x Genomics Xenium pipeline (https://www.10xgenomics.com/analysis-guides/performing-3d-nucleus-segmentation-with-cellpose-and-generating-a-feature-cell-matrix) to assign each puncta to a nuclei. For *STBD1*, the density of puncta was too low to restrict analysis to solely the nuclei. Instead, the density of puncta at each voxel was computed to create a density map for *STBD1*, and high-density regions were counted as *STBD1*^+^ cells.

### 3.9 Data and Code Availability

scRNA-seq data clustering, integration and visualization was performed in R v4.2.1. SAMap was run in Python v3.9.19. R Markdown and Jupyter notebooks required to reproduce the analyses presented here are available on our GitHub page (https://github.com/shekharlab/DoubleCones).

## 3.10 Acknowledgements

This work was supported by the NIH grant R00EY028625 (K.S.), the McKnight Foundation (K.S.), and funds from the University of California, Berkeley (K.S., D.T.). K.S. acknowledges support from the Glaucoma Research Foundation and the Melza M. and Frank Theodore Barr Foundation. We thank Prof. Leo Peichl for helpful discussions about the origin of rods and related literature. We thank Mr. Vishruth Dinesh for helpful discussions on integrating photoreceptors using Seurat, and to Dr. Joshua Hahn for help with the lizard, opossum, squirrel, and human snRNA-seq data. We are grateful to Dr. Pin Lyu, Prof. Seth Blackshaw, Dr. Wenjun Yan, Prof. Joshua R. Sanes, Dr. Yohey Ogawa, and Prof. Joseph Corbo for helpful advice and assistance in accessing their published datasets.

## 3.12 Author Contributions

D.T.: Conceptualization, Analysis, Software, Writing; T.Y.: Experiment, Analysis, Writing, T.B.: Conceptualization, Analysis, Writing; K.S.: Conceptualization, Analysis, Funding Acquisition, Writing.

---

**Figure S1:**
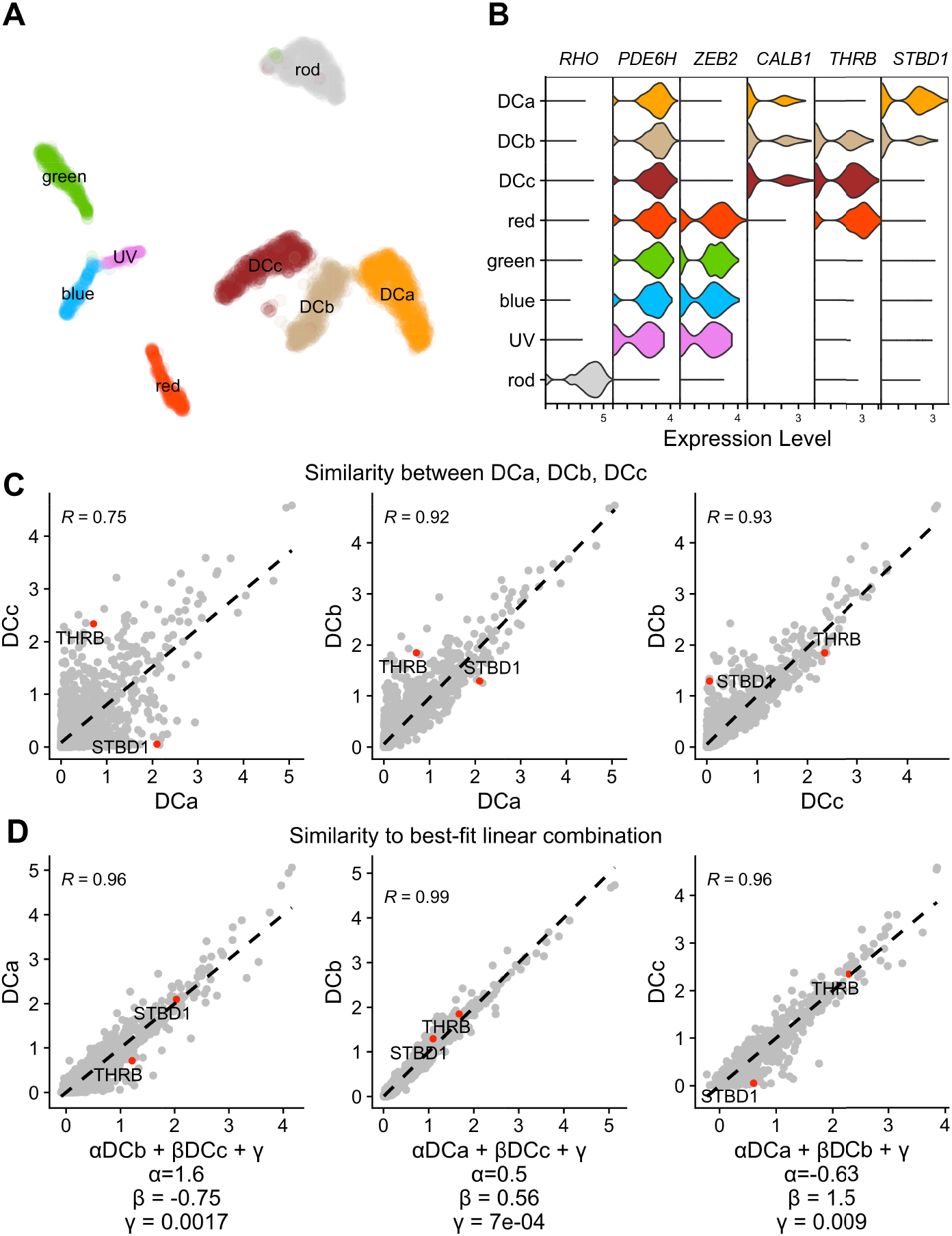
Identification of principal, accessory, and full double cones in the chicken retinal atlas. A) 2-D UMAP embedding of chicken retinal atlas ^17^. Three double cone (DC) clusters annotated by the authors, labeled DCa, DCb, and DCc, are highlighted. B) Violin plot showing the normalized and log-transformed expression values for marker genes in the photoreceptor clusters in panel A. Shown are the rod marker *RHO*, the cone marker *PDE6H*, the ancestral cone marker *ZEB2*, the double cone marker *CALB1*, the red cone marker *THRB*, and a novel marker for DCa (*STBD1*). Notice that DCb expresses *THRB* and *STBD1* at intermediate levels compared to DCa and DCc. C) Pairwise gene expression correlations between DCa, DCb, and DCc. DCb is similar to both DCa and DCc. Gene expression correlation of DCa, DCb, and DCc to the best-fit linear combination of the other two clusters. Middle panel shows that DCb is an average of DCa and DCc, indicating that it represents full, intact double cones (both principal and accessory member) entering a single 10x droplet.

**Figure S2:**
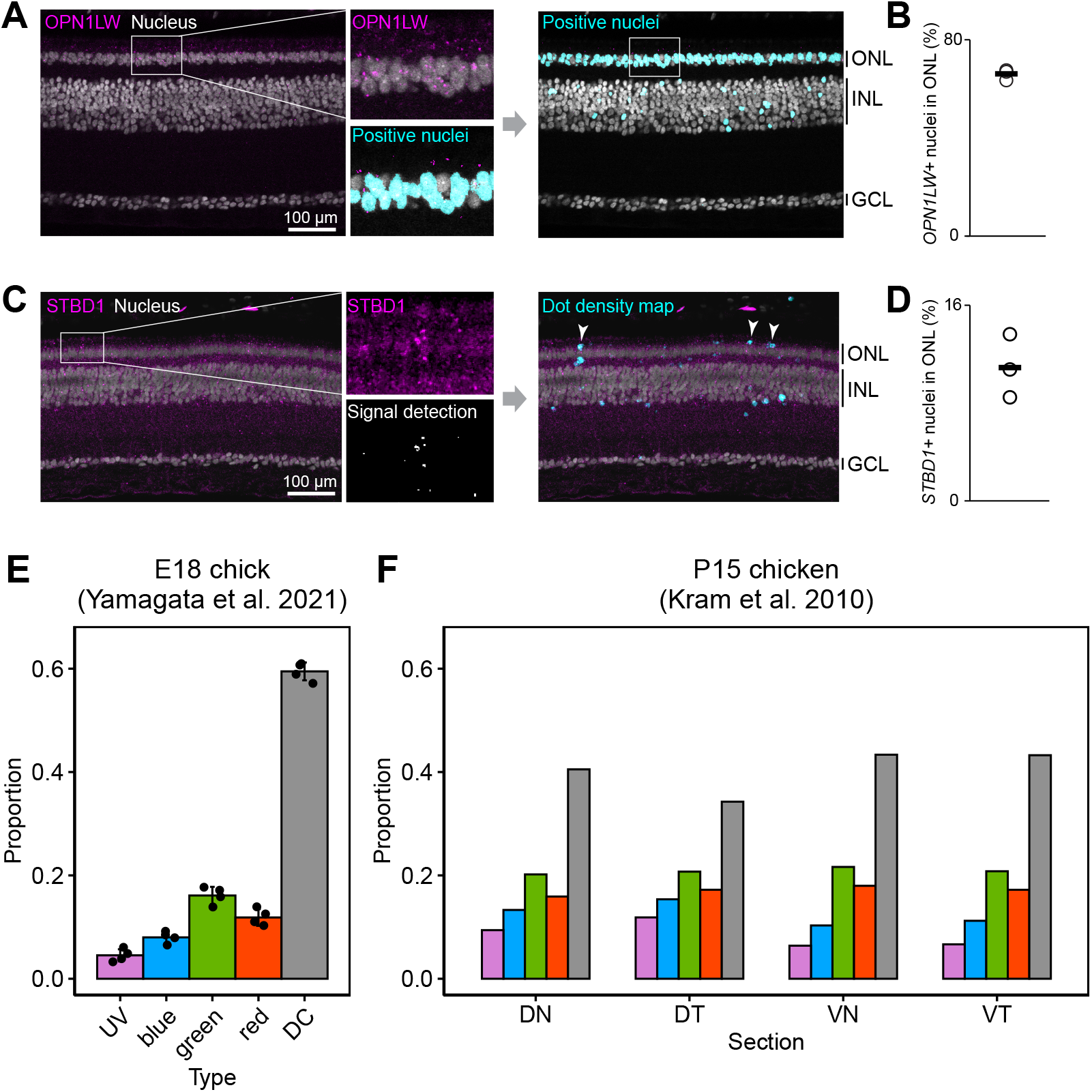
Analysis of photoreceptor proportions in anole lizard and chicken. A) *In situ* Hybridization Chain Reaction (HCR) targeting *OPN1LW* in retinal cross sections of the anole lizard (left panel). RNA signal and nuclei (Hoechst) are represented in magenta and grey, respectively. In the inset (middle), nuclei positive for *OPN1LW* are highlighted in cyan (details in **Methods**). Right panel shows *OPN1LW+* nuclei in the same field of view as the left panel. ONL, outer nucleus layer; INL, inner nucleus layer; GCL, ganglion cell layer. B) Percentage of *OPN1LW+* nuclei in the ONL. Each circle is a biological replicate. A total of 646 nuclei were counted in the ONL and the typical field of view was 640×540 μm. C) Same as A, but for HCR experiments targeting *STBD1*. Regions of high HCR signal density are shown in cyan. D) Percentage of *STBD1+* nuclei in the ONL. Each circle is a biological replicate. A total of 407 nuclei were counted in the ONL and the typical field of view was 640×380 μm. E) Relative proportions of photoreceptor types from scRNA-seq of E18 chicken retina ^17^. In these calculations, we estimated the total number of double cones to be the sum of the number of intact double cones (DCb) and the average of the numbers of the principal (DCc) and accessory (DCa) cells. F) Relative proportions of photoreceptor types from immunostaining of P15 chicken (Figure 2B of Kram et al. 2010^7^, reproduced using automeris.io WebPlotDigitizer). Photoreceptor types are colored the same as in panel C. The x-axis shows the proportions for tissue sections obtained from different quadrants of the retina: dorsonasal (DN), dorsotemporal (DT), ventronasal (VN), ventrotemporal (VT). Differences in proportions between panels C and D may be due to biases in cell capture in scRNA-seq or due to differences in age (E18 vs. P15), or both.

**Figure S3:**
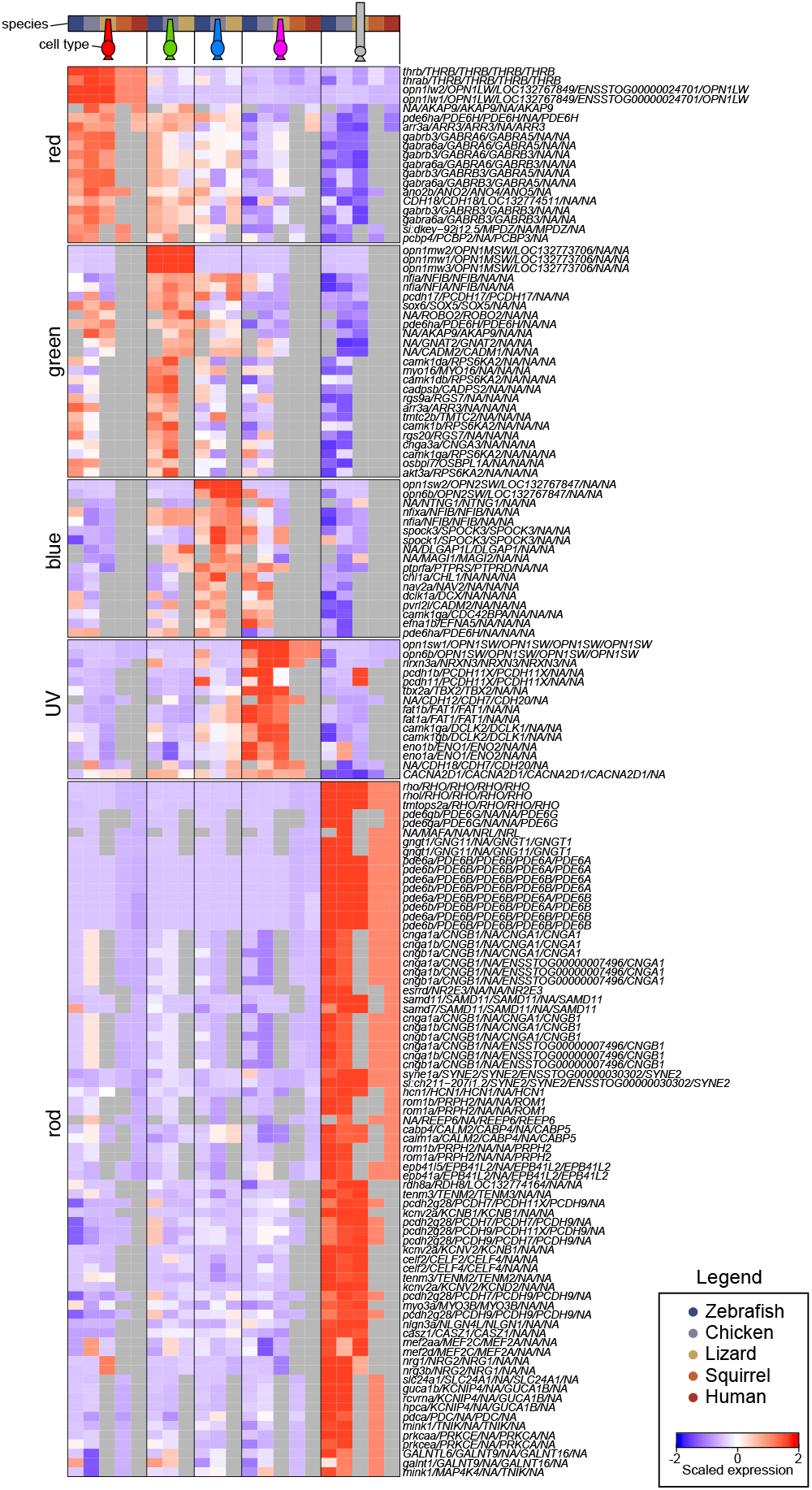
Conserved transcriptional signatures for rods and single cones. Heatmap of scaled gene expression showing the top conserved transcriptional signatures of ancestral photoreceptors (same as **Figure 2D**, but with gene names shown). Gene names are in the same order as the species (zebrafish, chicken, lizard, squirrel, and human). SAMap identified known paralogous relationships; for example, *opn1mw1/2/3* in zebrafish. Missing genes are indicated as *NA*.

**Figure S4:**
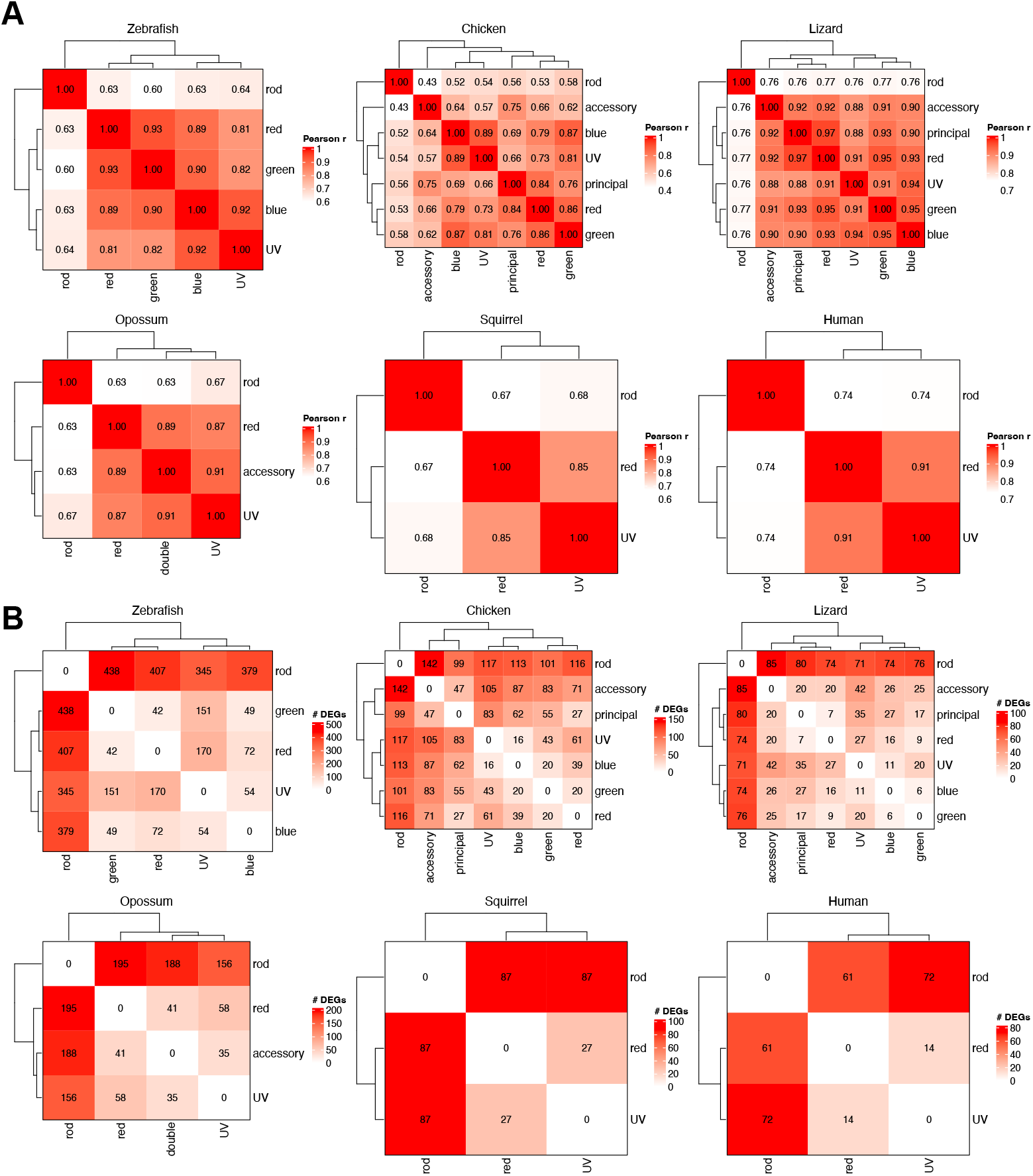
Pairwise gene expression similarity between photoreceptor types. A) Pearson correlation between average expression levels of photoreceptor clusters within species. Normalized and log-transformed counts were used to calculate correlations. B) Heatmaps showing number of differentially expressed genes (Benjamini-Hochberg-adjusted p *<* 0.001 and log fold change *>* 1) between clusters for each species.

**Figure S5:**
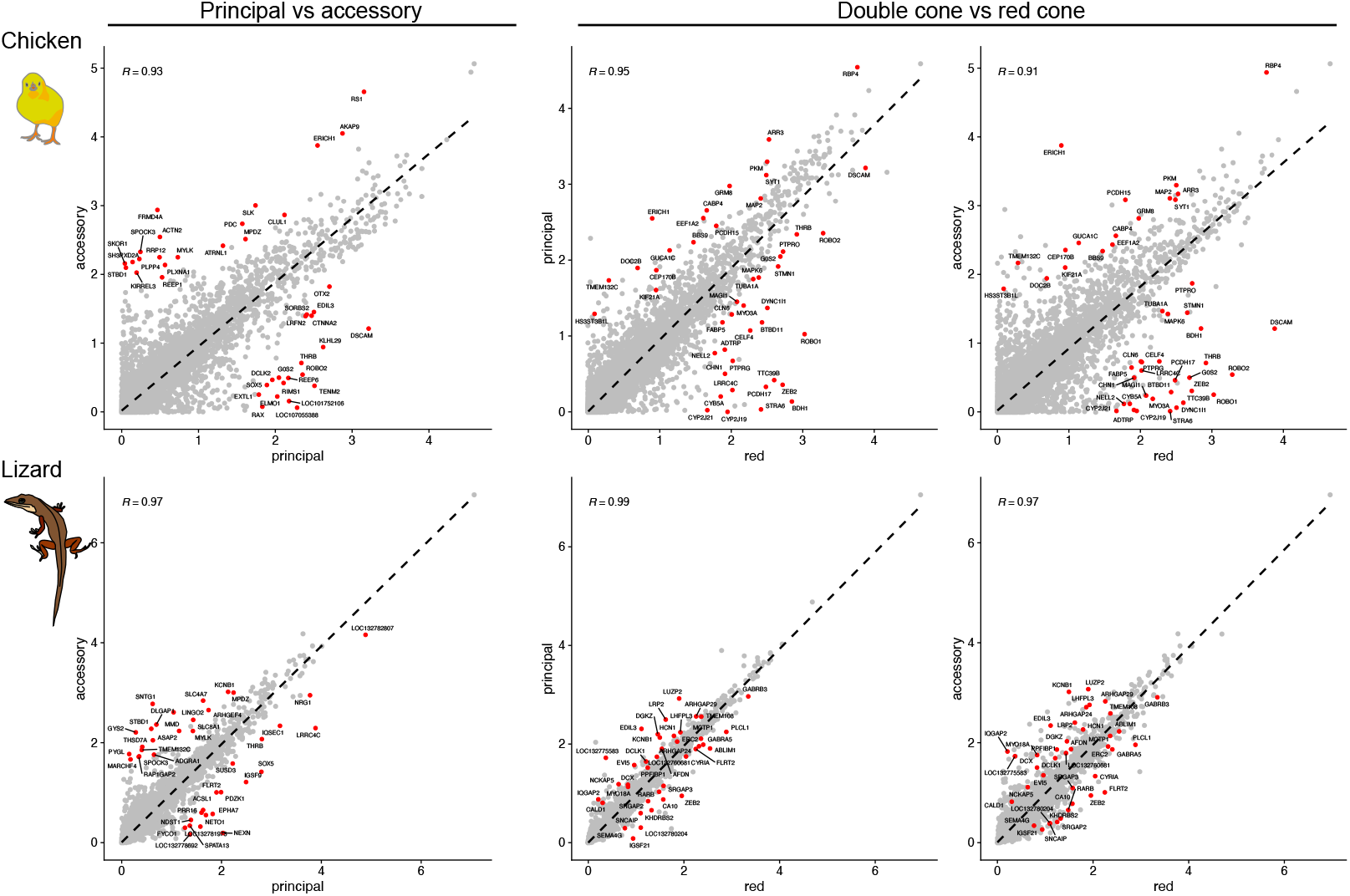
Genes that distinguish the principal from the accessory and the double cones from the red cone. Scatterplots comparing gene expression (normalized and log-transformed) between the principal and the accessory (left) or the two double cone members and the red cone (right). In cases where too many DEGs were returned, only the top DEGs were labeled – a full list is provided in **Table S2**. The top row shows the results for chicken, and the bottom panel shows the results for lizard. Note that Pearson correlation coefficients differ from **Figure S4** because only highly variable genes were considered in **Figure S4**, whereas all genes were considered for DE analysis.

**Figure S6:**
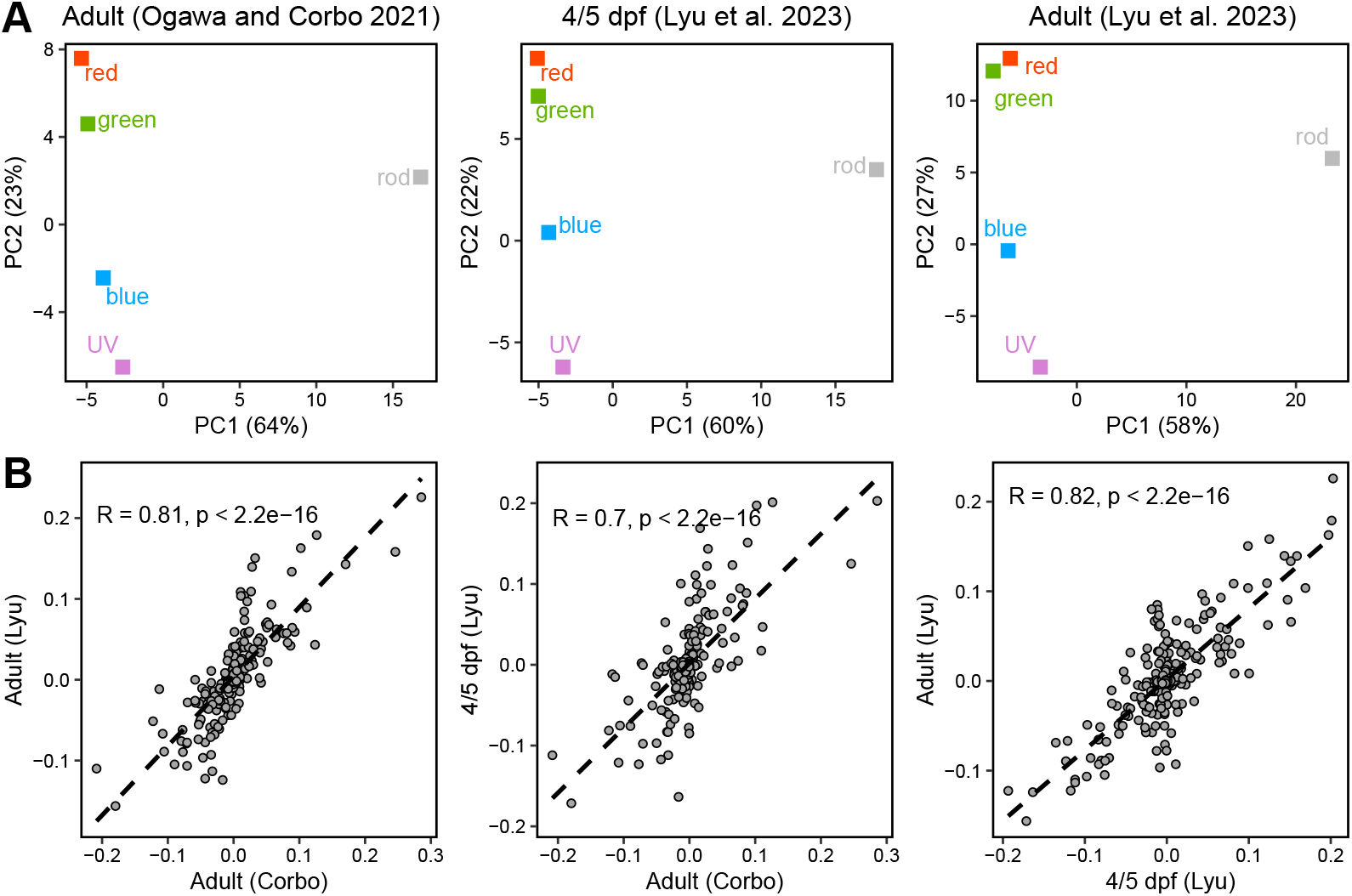
PC2 in zebrafish is reproducible in two different datasets. A) PCA of averaged photoreceptor gene expression profiles within three zebrafish datasets (columns). In all three cases, *PC*1 separates rods and cones, while *PC*2 separates cone types based on their color. B) Scatterplots comparing *PC*2 gene loadings between pairs of datasets in panel A.

**Figure S7:**
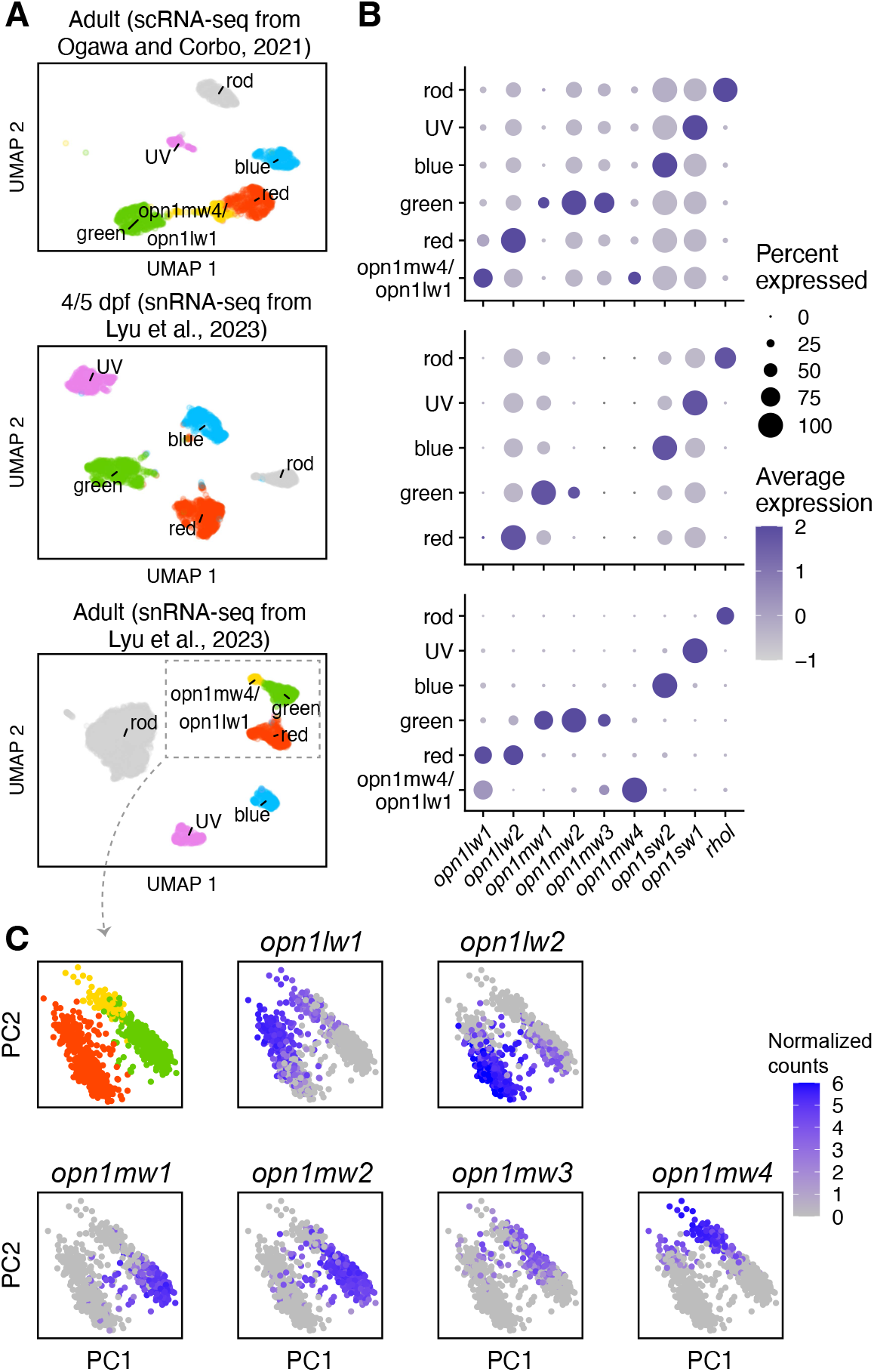
The *opn1mw4/opn1lw1+* cones first described in Ogawa and Corbo 2021 are present in adult zebrafish but not in the developing zebrafish. A) UMAP embeddings for three zebrafish photoreceptor atlases:1) adult scRNA-seq (Ogawa and Corbo 2021), 2) 4/5 days post-fertilization (dpf) snRNA-seq (Lyu et al. 2023), and 3) adult snRNA-seq (Lyu et al. 2023). B) Dotplots showing the expression of the visual opsins. The yellow cluster expresses both *opn1mw4* and *opn1lw1* and is adult-specific, suggesting it develops after 4/5 dpf. C) PCA of reclustered red, green, and *opn1mw4/opn1lw1+* cones from Lyu et al. adult data. The top left panel is colored by identity like in panel B. The rest of the panels are colored by the expression of the various red and green opsins. Notice that both the red and green opsins have a graded expression pattern, suggesting the presence of specialized subsets that may be enriched in some retinal regions.

